# Microfluidic analysis reveals ROCK2 regulation of endothelial cilia is essential for blood vessel lumen formation and vascular integrity

**DOI:** 10.64898/2026.05.27.728336

**Authors:** S. Zahed Mohajerani, G. Grant, A. McCarthy, M.D. Bourn, Peyman, C.A. Johnson, G. Mavria

## Abstract

**Background:** The formation of a patent vascular lumen is fundamental to circulatory function, a process governed by cytoskeletal dynamics and mechanosensory signalling. Endothelial cilia are present during blood vessel lumen development, but their precise functional role remains poorly understood.

Understanding how cilia coordinate with endothelial cytoskeletal and signalling pathways is critical for elucidating mechanisms of vascular morphogenesis.

**Methods:** We have established a microfluidic system that recapitulates endothelial tube formation under fluid flow, enabling pharmacological and genetic manipulation with real-time visualisation of tube behaviour. Cilia, cytoskeletal dynamics, and lumen development were analysed *in vitro*, and *in vivo*.

**Results:** Early perfusion in the microfluidic system induced a hierarchical vascular network. Inhibiting Rho-kinase (ROCK) or knocking down ciliary components (IFT88 and RPGRIPL1) suppressed lumen formation. ROCK inhibition or genetic ablation disrupted cilia in endothelial and non-endothelial cells, associated with LIM-kinase inhibition. Crucially, ROCK2 genetic ablation caused endothelial cilia loss, misorientation, and abrogated lumen formation, leading to haemorrhages and compromised vascular integrity in vivo.

**Conclusions:** Our findings unveil a previously unrecognised co-regulation between cilia and ROCK signalling essential in vascular lumen formation.

## Introduction

The formation of a patent vascular lumen is fundamental to establishing a circulatory system capable of delivering oxygen and nutrients to tissues. The intricate process of lumen formation transforms endothelial cords into hollow, perfusable tubes, a complex morphogenetic event governed by the precise integration of genetic programming, cytoskeletal dynamics, and mechanosensory signalling.^1,2^

Following initial vasculogenesis and angiogenic sprouting, nascent vessels establish a lumen through cord hollowing, cell hollowing, or membrane invagination, processes that require endothelial cell polarisation, junctional reorganisation, and actomyosin-driven cell shape changes.^3,4^ Subsequent lumen expansion and network maturation are critically dependent on haemodynamic forces, with blood flow and pressure providing mechanical cues that drive inverse blebbing and the formation of a stable, size-appropriate conduit.^4^ Beyond lumen expansion, flow forces also induce endothelial cell (EC) polarisation, with the Golgi apparatus localising upstream of the nucleus in response to shear stress, a process that directs migration against flow during vascular remodelling.^5,6^

The actin cytoskeleton is a central effector of these morphological changes, with actin dynamics tightly controlled by the Rho family of GTPases and their downstream effectors, including the Rho-associated kinases ROCK1 and ROCK2.^7,8^ ROCK signalling, through phosphorylation of targets such as myosin light chain (MLC) and LIM-kinase (LIMK), regulates actomyosin contractility and stress fibre formation, and consequently influences EC adhesion, migration, and tube morphology.^9^ While ROCK has been implicated in lumenogenesis, its precise role is complex and stage-dependent, with evidence suggesting it is dispensable for initial lumen opening but crucial for later diameter regulation and stability.^8^ Although shear stress is a primary driver of vascular growth and remodelling, pressure-derived circumferential stretch also plays a critical role in regulating vessel calibre and EC behaviour.^10,11^ Distinguishing the relative contributions of these forces remains an active area of investigation.

A key unresolved question is how ECs sense and transduce the physical force of blood flow into the cytoskeletal remodelling required for lumen formation. The primary cilium, a solitary, microtubule-based organelle projecting from the apical surface of quiescent ECs, is a prime candidate for this mechanosensory role.^12,13^ Cilia function as cellular antennae, housing mechanosensitive channels such as polycystin-1 (PKD1) and polycystin-2 (PKD2), and they are essential for flow-sensitive calcium signalling and nitric oxide release in vitro.^12,14^ In vivo, ciliary dysfunction is linked to vascular pathologies, including haemorrhage and defective angiogenesis.^15^ Intriguingly, the assembly, maintenance, and disassembly of cilia are regulated by the actin cytoskeleton and its upstream regulators.^16,17^ Actin polymerisation generally inhibits ciliogenesis, while actin depolymerisation promotes it.^16^ Given that Rho/ROCK signalling is a master regulator of actin dynamics, a potential link between ROCK activity and ciliary function in ECs exists but remains largely unexplored.

Studying the interplay of flow, cytoskeletal dynamics, and ciliary signalling in vivo is challenging due to technical limitations and system complexity. In vitro models, particularly three-dimensional co-culture systems, recapitulate key steps of angiogenesis and tubulogenesis^18^. However, traditional static cultures lack the essential haemodynamic component. Microfluidic organ-on-chip platforms address this gap by enabling precise control of fluid flow and shear stress, offering a biomimetic environment to study vascular morphogenesis and function in real time. In this study, we employ a perfused microfluidic organotypic co-culture model to investigate the role of EC cilia and ROCK signalling in blood vessel lumen development. We demonstrate that fluid flow drives the formation of a hierarchical vascular network. Using this platform, combined with pharmacological and genetic interventions in vitro and in vivo, we reveal that functional cilia, regulated by ROCK2 signalling, are essential for proper lumen formation and stability. Our findings uncover a novel mechanistic co-regulation between the mechanosensory cilium and a key cytoskeletal regulator during vascular morphogenesis.

## Materials and Methods

### Cell culture

The following cell lines were cultured in a humidified atmosphere at 37°C with 5% CO_2_. hCMEC/D3 (human cerebral microvascular endothelial cells) (ATCC® CRL-3245™) were cultured in EndoGRO-MV Complete Media (Merck, SCME004) supplemented with 1 ng/ml basic Fibroblast Growth Factor (bFGF; PeproTech, 100-18B) and used between passages 30-35 for ciliogenesis assays. bEnd.3 (mouse brain endothelial cells) (ATCC® CRL-2299™) were cultured in DMEM (Gibco, 11995065) supplemented with 10% (v/v) heat-inactivated foetal bovine serum (FBS) (Merck, F9665), 100 U/ml penicillin, 100 µg/ml streptomycin, and 2 mM L-glutamine (all from Gibco). hTERT RPE-1 (human retinal pigment epithelial cells) (ATCC® CRL-4000™) were cultured in DMEM/F12 (Gibco, 11330032) supplemented with 10% FBS, penicillin/streptomycin, and L-glutamine. Cells between passages 28-35 were used for ciliogenesis assays. Human Dermal Fibroblasts (HDFs) (Cellworks, ZHF-C) were cultured in DMEM supplemented with 10% FBS, penicillin/streptomycin, and L-glutamine, and used between passages 8-11. Human Umbilical Vein Endothelial Cells (HUVECs) (Cellworks, ZHC-3110) were cultured in Endothelial Cell Growth Medium-2 (EGM-2, Lonza, CC-3162) or Human Large Vessel Endothelial Cell Growth Medium (Cellworks, ZHC-4101) with supplied supplements. HUVECs were used for experiments between passages 1-4. HEK-293T (human embryonic kidney cells) (ATCC® CRL-3216™) were cultured in DMEM supplemented with 10% FBS, penicillin/streptomycin, and L-glutamine, and used for lentiviral production until passage 11. For ciliogenesis assays, cells were seeded onto 35 mm glass-bottom dishes (Ibidi) and grown to ∼70% confluence. To induce quiescence and cilia formation, cells were serum-starved (0.5% FBS) for 18 hours, then fixed and immunostained.

### Antibodies and Reagents

The following primary antibodies were used: rabbit anti-ARL13B (Proteintech, 17711-1-AP, 1:250), rat anti-endomucin (Santa Cruz, sc-65495, 1:100), rat anti-CD31 (Santa Cruz, sc-31045, 1:100), rabbit anti-ROCK2 (Bethyl, A300-046A, 1:200), rabbit anti-NG2 (Millipore, AB5320, 1:200), rabbit anti-IFT88 (Proteintech, 13967-1-AP, 1:1000), rabbit anti-RPGRIP1L (Proteintech, 55160-1-AP, 1:1000), mouse anti-β-actin (Sigma-Aldrich, A6441, 1:5000), and rabbit anti-GAPDH (Abcam, ab9485, 1:1000). Secondary antibodies were Alexa Fluor 488-or 594-conjugated goat anti-mouse, anti-rabbit, or anti-rat IgG (Sigma-Aldrich, 1:500), horseradish peroxidase (HRP)-conjugated goat anti-rat IgG (Vector Laboratories, PI-9400, 1:500), HRP-conjugated goat anti-rabbit IgG (Vector Laboratories, PI-1000, 1:500), HRP-conjugated anti-rabbit IgG (Cell Signaling Technology, 7074S, 1:5000), and HRP-conjugated anti-mouse IgG (Cell Signaling Technology, 7076S, 1:5000). The following reagents were used: DAPI (Sigma-Aldrich, D9542), BMS-5 (Enzo, ENZ-CHM178), Latrunculin B (Sigma-Aldrich, L5288), and Y27632 (Tocris, 1254).

### Lentivirus Production and Transduction

#### Plasmid preparation

shRNA plasmids were transformed into competent *E. coli* and cultured in LB broth with 100 µg/ml ampicillin at 37°C. Plasmid DNA was purified using the PureLink HiPure Plasmid Midiprep Kit (Invitrogen) following the manufacturer’s protocol.

#### shRNA constructs

Doxycycline-inducible shRNA constructs targeting human IFT88 (Gene ID: 8100) and RPGRIP1L (Gene ID: 23322) were obtained from Horizon Discovery (formerly Open Biosystems) as pTRIPZ lentiviral vectors. The pTRIPZ vector contains a doxycycline-inducible TRE3G promoter driving shRNA expression and a TurboRFP reporter to monitor transduction efficiency. A non-targeting shRNA construct (Horizon Discovery, RHS4743) was used as a negative control.

#### Lentivirus production

Lentiviral vectors were produced in HEK-293T cells using calcium phosphate transfection. Briefly, for each 10 cm dish, a DNA precipitate containing 20 µg transfer vector (pTRIPZ shRNA), 15 µg psPAX2 packaging plasmid, and 6 µg pMD2.G envelope plasmid was formed in 0.5 mL ddH_2_O with 50 µL 2.5 M CaCl_2_, then mixed with 0.5 mL 2× HEPES-buffered saline. The precipitate was added to cells. Viral supernatant was collected at 48 and 72 hours post-transfection, pooled, and filtered through a 0.45 µm filter.

#### Transduction of HUVEC

HUVEC were plated in T75 flasks and grown to approximately 70% confluence. For transduction, 5 × 10^5^ HUVECs were incubated with viral supernatant plus 8 µg/ml polybrene (Sigma-Aldrich, TR1003G) for 16 hours. Cells were transduced with pTRIPZ vectors expressing TurboRFP or doxycycline-inducible shRNAs targeting IFT88, RPGRIP1L, or a non-targeting control.

#### Doxycycline induction

Knockdown was induced by adding 2 µg/ml doxycycline (Sigma-Aldrich, D9891) to the culture medium on day 9 of the co-culture assay. Media were replenished with fresh doxycycline on days 10, 12, and 13. Knockdown efficiency was confirmed by immunoblotting as described above.

### Western blotting

Cells were washed twice with ice-cold PBS and lysed in RIPA buffer (50 mM Tris-HCl pH 7.4, 150 mM NaCl, 1% NP-40, 0.5% sodium deoxycholate, 0.1% SDS) supplemented with protease inhibitor cocktail (Roche, 11836170001) and phosphatase inhibitor cocktail (Thermo Scientific, 78420). Lysates were incubated on ice for 20 minutes with occasional vortexing, then centrifuged at 14,000 × g for 15 minutes at 4°C. The supernatant was collected, and protein concentration was determined using a BCA protein assay kit (Thermo Scientific, 23225). Equal amounts of protein (20 µg per lane) were mixed with Laemmli sample buffer (Bio-Rad, 1610747) containing 5% β-mercaptoethanol, boiled at 95°C for 5 minutes, and resolved by SDS-PAGE on 10–15% Tris-Glycine gels (Bio-Rad, 5678084). Proteins were transferred onto nitrocellulose membranes (Hybond, GE Healthcare, 10600033) using a wet transfer apparatus (Bio-Rad, Mini Trans-Blot Cell) at 100 V for 1 hour at 4°C in transfer buffer (25 mM Tris, 192 mM glycine, 20% methanol). Membranes were blocked in 5% non-fat dry milk (Bio-Rad, 1706404) or 5% bovine serum albumin (BSA; Sigma-Aldrich, A9418) in Tris-buffered saline containing 0.1% Tween-20 (TBST) for 1 hour at room temperature. Membranes were then incubated with primary antibodies (see Antibodies and Reagents) diluted in blocking buffer. After washing, membranes were incubated with HRP-conjugated secondary antibodies (see Antibodies and Reagents) diluted 1:5000 in blocking buffer for 1 hour at room temperature. Immunoblots were visualised using enhanced chemiluminescence (ECL) substrate (Thermo Scientific, 34580) on a Bio-Rad ChemiDoc imaging system (Bio-Rad, ChemiDoc MP). Images were processed using Image Lab software (Bio-Rad, version 6.1).

### Immunofluorescence

Cultured cells, prior to fixation, cells were rinsed with PBS and then fixed in 4% paraformaldehyde (PFA) for 20 minutes at room temperature (RT). Cells were permeabilised with 0.1% Triton X-100 in PBS for 10 minutes at RT and blocked in 0.5% BSA/PBS for 1 hour at RT. Primary antibodies (see Antibodies and Reagents) were diluted in blocking buffer and incubated overnight at 4°C. After washing three times with PBS (5 minutes each), cells were incubated with Alexa Fluor-conjugated secondary antibodies (see Antibodies and Reagents) diluted in blocking buffer for 1 hour at RT. Nuclei were stained with DAPI (1 µg/ml) for 5 minutes at RT. Coverslips were mounted using Fluorescent Mounting Medium (Dako, S3023). Images were acquired using a Nikon A1R confocal microscope (Nikon Instruments) with a 40× objective (NA 0.95).

Mouse embryos were harvested at E13.5, fixed in 4% PFA for 30 minutes at RT, and processed as whole mounts. Samples were permeabilised with 0.1% Triton X-100 in PBS for 1 hour at RT, blocked with 5% BSA/PBS for 1 hour at RT, and incubated with primary antibodies (see Antibodies and Reagents) diluted in blocking buffer overnight at 4°C. After washing three times with PBST (30 minutes each), samples were incubated with Alexa Fluor-conjugated secondary antibodies (see Antibodies and Reagents) diluted in blocking buffer for 2 hours at RT. Nuclei were stained with DAPI (1 µg/ml) for 10 minutes at RT. Z-stack images were acquired at 1 µm intervals using a 40× oil immersion lens (NA 1.3) on a Nikon A1R confocal microscope. To assess vascular development, the superficial pial plexus was visualised in whole-mount telencephalon preparations at E13.5. Four embryos were analysed, with five fields of view quantified per embryo.

### Immunohistochemistry

E13.5 embryos were genotyped, fixed in 4% paraformaldehyde (PFA) in phosphate-buffered saline (PBS) for 24 hours at 4°C, then processed for paraffin embedding. Embedded tissues were sectioned at 5 µm thickness using a rotary microtome (Leica RM2235). Sections were mounted on SuperFrost Plus glass slides (Thermo Fisher Scientific, J1800AMNZ). Sections were deparaffinised in xylene (3 × 5 minutes) and rehydrated through a graded ethanol series (100%, 95%, 70%, 50%; 3 minutes each) followed by distilled water. Antigen retrieval was performed by heating sections in 10 mM sodium citrate buffer (pH 6.0) containing 0.05% Tween-20 using a pressure cooker at 120°C for 15 minutes. Sections were allowed to cool to room temperature for 20 minutes and then washed in PBS containing 0.1% Tween-20 (PBST). Endogenous peroxidase activity was quenched by incubating sections in 3% hydrogen peroxide (H_2_O_2_) in methanol for 15 minutes at room temperature. Sections were then blocked in 5% normal goat serum diluted in PBST for 1 hour at room temperature. Primary antibodies (see Antibodies and Reagents) were diluted in blocking buffer and incubated overnight at 4°C in a humidified chamber. After washing three times with PBST (5 minutes each), sections were incubated with HRP-conjugated secondary antibodies (see Antibodies and Reagents) diluted 1:500 in blocking buffer for 1 hour at room temperature. Antibody binding was visualised using 3,3’-diaminobenzidine (DAB) substrate (Vector Laboratories, SK-4100) according to the manufacturer’s instructions. Sections were counterstained with haematoxylin (Sigma-Aldrich, GHS316) for 30 seconds, dehydrated through graded ethanol (50%, 70%, 95%, 100%; 3 minutes each), cleared in xylene, and mounted with DPX mounting medium (Sigma-Aldrich, 06522). Brightfield images were captured using a Nikon Axiovision microscope (Nikon Instruments) with a ×40 objective (NA 0.75). For lumen analysis, transverse brain sections at the level of the interventricular foramen of Monro were used. All endomucin-positive lumens within the brain parenchyma were measured using the circle tool in Axiovision software (version 4.9). At least three sections per embryo from three embryos per genotype were analysed.

### Microfluidic Device Design and Fabrication

Single-chamber microfluidic devices were fabricated in polydimethylsiloxane (PDMS) based on a previously described design.^19^ The 5-pillar design was chosen as it recapitulates key aspects of early vascular development, including formation of capillary-like tubes, as opposed to the 3-pillar configuration which are more representative of precapillary arterioles.^20^ For fabrication, a master mold was created by patterning SU-8 2050 photoresist (Kayaku Advanced Materials) on a 4-inch silicon wafer using standard photolithography. The SU-8 was spin-coated to a thickness of 100 µm and baked according to the manufacturer’s protocol. PDMS base and curing agent (Sylgard 184, Dow Corning) were mixed at a 10:1 (w/w) ratio, degassed in a vacuum desiccator for 30 minutes, poured onto the master mold, and cured at 80°C for 2 hours. The cured PDMS was carefully peeled from the mold, and 1 mm (cell loading) and 2 mm (media reservoir) ports were punched using biopsy punches (Kai Medical, BPP-20F). Devices and glass coverslips (24 × 50 mm, No. 1.5; Thermo Fisher Scientific) were cleaned with isopropanol and dried with nitrogen gas. Both surfaces were activated by oxygen plasma (Harrick Plasma, PDC-002) at 30 W for 45 seconds and immediately bonded together to form an irreversible seal. Bonded devices were baked at 80°C for 10 minutes to strengthen the seal.

### Microfluidic Co-Culture Angiogenesis Assay

All cultures were maintained at 37°C in a humidified atmosphere with 5% CO_2_. Sterilised microfluidic devices were exposed to UV light for 30 minutes prior to cell loading. A co-culture hydrogel was prepared by mixing 6 × 10^4^ HUVECs and 6 × 10^4^ HDFs in 8 µL of 5 mg/ml fibrinogen (Merck, F3879) dissolved in sterile PBS, followed by addition of 0.6 µL of 50 U/ml thrombin (Merck, T6884). The cell-fibrinogen-thrombin mixture was immediately injected into the central chamber via the cell loading port and allowed to polymerise at 37°C for 20 minutes. After polymerisation, media reservoirs were filled with 2.5 mL of medium (50:50 DMEM:Large Vessel Endothelial cell growth medium (LVE; Cellworks, ZHC-4101)) supplemented with 25 ng/ml VEGFA (Merck, V7259) and 10 ng/ml bFGF (PeproTech, 100-18B) to establish hydrostatic pressure-driven flow (see Fig. 2A). Flow direction was reversed daily, and media were changed every 24 hours. On day 6, an additional 3 × 10^4^ HUVECs in 30 µL of LVE medium were injected into the side channels to promote anastomosis with developing tubules. From day 11, the medium was replaced with Angiogenesis Growth Medium (Cellworks, ZHA-3100). Phase-contrast and fluorescence images were acquired using an EVOS microscope (Thermo Fisher Scientific) on day 14.

### Mice, Embryos, and Timed Matings

All animal procedures were approved by the Animal Ethics Committees of the Institute of Cancer Research and the University of Leeds in accordance with National Home Office regulations under the Animals (Scientific Procedures) Act 1986. *Rock2* conditional knockout mice (*Rock2*f/f; exons 5-6 floxed) generated by Kümper and colleagues^21^ in the laboratory of the late Professor Christopher J Marshall were crossed to a transgenic line expressing improved Cre (icre) ubiquitously to generate global Rock2 knockout mice (*Rock2*^-/-^). Timed matings were set up between *Rock2*^+/-^ heterozygotes, and embryos were harvested at embryonic day (E) 13.5 for analysis. Male and female embryos were included, as sex cannot be distinguished at this developmental stage. All animals were maintained on a C57BL/6J background. Genotyping for the *Rock2* allele was performed by Transnetyx using real-time PCR. Littermate *Rock2*^+/+^ embryos were used as wild-type controls. Randomization was not applicable as genotype was determined by Mendelian inheritance from heterozygote intercrosses, and all embryos of the appropriate developmental stage were harvested regardless of genotype. Statistical analysis is described in the Statistical analysis section below.

### Statistical analysis

Normal distribution of data was assessed before parametric tests were carried out. Statistical significance was determined using a two-tailed Student’s t-test assuming unequal variance between sample means, performed using GraphPad Prism (version 8; GraphPad Software, Boston, MA, USA). Pairwise comparisons were performed where appropriate. Data are presented as mean ± standard error of the mean (SEM) from at least three independent biological replicates. A p-value ≤ 0.05 was considered statistically significant.

## Results

### Endothelial cells extend primary cilia during active vascular lumen development

While primary cilia have been documented in the retinal vasculature^22^ their prevalence, distribution, and functional role during active vascular lumen formation remain poorly characterised. These sensory organelles undergo dynamic length regulation, a process governed by intraflagellar transport (IFT) and coordinated with the cell cycle^23^ (Fig. 1A). We systematically investigated the presence of endothelial primary cilia across multiple *in vitro* and *in vivo* models.

**Figure 1.**
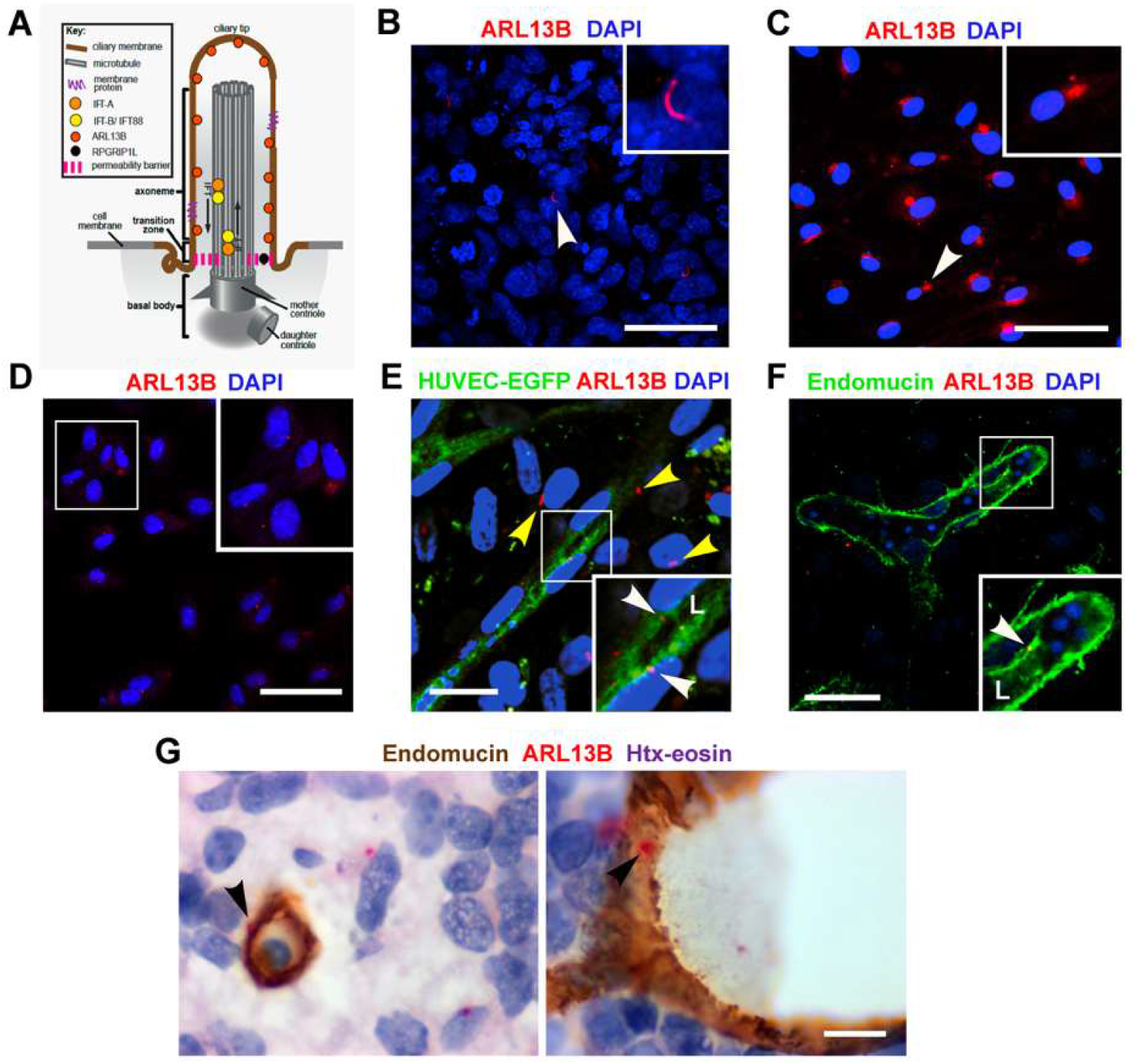
Endothelial primary cilia are present across multiple models of vascular development. **(A)** Schematic of a primary cilium, depicting its core compartments (basal body, transition zone, axoneme, ciliary tip) and the localization of key ciliary proteins ARL13B and RPGRIP1L. Intraflagellar transport (IFT) complexes (IFT-A and IFT-B) facilitate cargo trafficking along the axoneme. Adapted from Smith et al. (2020). **(B, C)** Immunofluorescence images confirming primary cilia (ARL13B, red) in human cerebral microvascular endothelial cells (hCMEC/D3) **(B)** and mouse bEnd.3 cells **(C)**. Nuclei are counterstained with DAPI (blue). Scale bars, 50 µm. **(D)** Primary cilia (ARL13B, red) in a monolayer of human umbilical vein endothelial cells (HUVECs). Nuclei are stained with DAPI (blue). Scale bar, 50 µm. **(E)** Primary cilia (ARL13B, red) on endothelial tubules (white arrowheads) and fibroblasts (yellow arrowheads) in a HUVEC–human dermal fibroblast (HDF) organotypic coculture. L, lumen. Nuclei are stained with DAPI (blue). Scale bar: 15 µm. **(F, G)** Endothelial primary cilia in E13.5 mouse embryonic brain vessels. Cilia (arrows, ARL13B, red) are visible on endothelial cells lining the lumen of a superficial vessel **(F)** and on intraparenchymal vessels **(G)**. Endomucin (green (F), brown (G)) marks endothelial cells; nuclei are stained with DAPI (blue). Scale bars, 50 µm **(F)** and 10 µm **(G)**.

Immunofluorescence analysis confirmed that primary cilia were present on human (hCMEC/D3) and mouse (bEnd.3) cerebral microvascular endothelial cells (Fig. 1B, C). This finding was conserved in human umbilical vein endothelial cells (HUVECs), both in standard two-dimensional monolayers (Fig. 1D) and in a three-dimensional organotypic co-culture system with fibroblasts, where cilia were observed projecting from endothelial cells into the forming tubes (Fig. 1E). To validate these observations in a physiological context, we examined embryonic mouse brains at E13.5, a key stage of active brain angiogenesis and lumen formation. We detected abundant primary cilia projecting from endothelial cells lining both superficial cortical blood vessels (Fig. 1F) and vessels embedded within the brain parenchyma (Fig. 1G). Collectively, these data demonstrate that primary cilia are a ubiquitous feature of endothelial cells from diverse vascular beds and are present during active vascular morphogenesis and lumen development.

### A microfluidic co-culture system reveals hierarchical vascular tube formation and ROCK-dependent lumenogenesis

Having established the presence of endothelial cilia in our 3D organotypic co-culture model (Fig. 1E), we sought to investigate their role under physiologically relevant flow conditions. We employed a microfluidic platform engineered to recapitulate *in vivo*-like endothelial tube formation (Fig. 2A; Fig. S1). In this system, HUVEC-EGFP and human dermal fibroblasts (HDF) were co-cultured in a fibrin gel within the microfluidic chamber, enabling real-time imaging and longitudinal assessment of vascular morphogenesis. This approach allowed lumen formation to be quantified in defined regions of the chamber (Fig. S1A), with lumenised length expressed as a percentage of total vessel length (Fig. S1B).

Under flow, the co-culture supported sprouting angiogenesis and progressive lumen development, culminating in the formation of an interconnected vascular network within 7 days co-coculture (Fig. 2B; Fig. S2). By day 14, the network had matured into a closed, perfusable system, with interconnected tubes spanning the central chamber and connecting to the device’s side channels (Fig. 2B). This configuration established interstitial flow across the matrix, followed by directional luminal flow within the developing tubes (Fig. S3). The establishment of perfusion drove the emergence of a hierarchical vascular architecture. We observed wide conduit vessels near the chamber periphery and narrow capillary-like tubes 10-15 µm in diameter in the central chamber (Fig. 2C). Larger tubes contained distinct endothelial pillars, suggestive of active intussusceptive angiogenesis (Fig. 2C).

**Figure 2.**
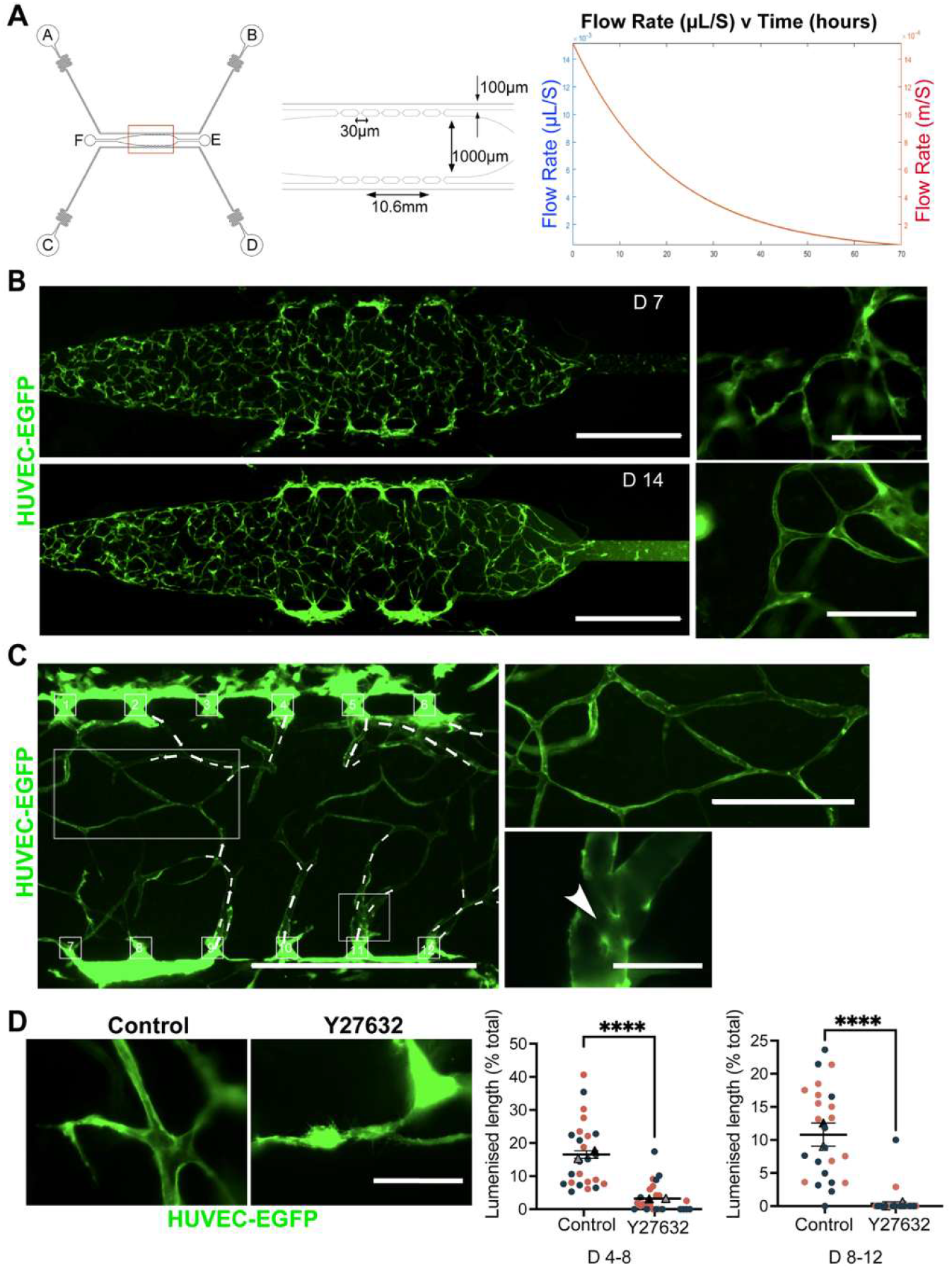
Microfluidic analysis of lumen development and requirement for ROCK signalling under conditions of fluid flow. **(A)** Schematic of the microfluidic device. A central chamber is connected to four reservoirs (A–D) via side channels. Cells in a fibrin gel are loaded through port E. The corresponding graph depicts flow-rate variation over time between reservoirs A–B and C–D, with a channel flow rate maintained at 50–60 µl/h. **(B)** Epifluorescence images showing progressive tubule and lumen formation in organotypic co-cultures at day 7 (partial network) and day 14 (complete network). Scale bars: 1000 µm (left panels); 200 µm (right panels). **(C)** Hierarchical vessel organisation under flow at day 14. White arrows indicate flow direction (left panel). Higher-magnification views show small-diameter (top) and large-diameter (bottom) tubes; arrowhead indicates a pillar structure within the lumen. Scale bars: left panel, 1000 µm; top right panel, 400 µm; bottom right panel, 100 µm. **(D)** Effect of ROCK inhibition (Y27632, 10 µM) on lumenisation between days 4–8. Scale bar: 100 µm. Dot plots quantify the percentage of lumenised length during days 4–8 and days 8–12. Data are presented as mean ± SEM from two independent experiments (n = 24 channel squares (explained in Fig. S1) per condition. Statistical significance was determined using an unpaired two-tailed Student’s t-test (****, p < 0.0001).

To determine the contribution of ROCK signalling to this process, we performed pharmacological inhibition at two distinct stages of vascular development: an early treatment window (days 4–8), prior to the onset of lumen formation, and a late treatment window (days 8–12), after tubes had established and begun to mature. ROCK inhibition significantly reduced lumen development in both treatment windows, disrupting tube morphology and decreasing the length of structures with lumens (Fig. 2D). These observations demonstrate that ROCK activity is required not only for initial lumen establishment but also for subsequent expansion and maintenance of structural integrity under flow.

### Primary cilia are required for endothelial lumen formation and expansion

Given the requirement of ROCK for lumenogenesis in the microfluidic system (Fig. 2D) and the known regulation of cilia by cytoskeletal pathways, we hypothesised that endothelial primary cilia are functionally required for tube formation. To test this, we used a doxycycline-inducible shRNA system in our microfluidic model to disrupt key ciliary components (Fig. S4). We first induced knockdown of the ciliary protein RPGRIP1L (Fig. 3A, B), which led to pronounced disruption of lumen formation (Fig. 3E). To confirm specificity, we additionally targeted IFT88, a core intraflagellar transport protein critical for cilium assembly and maintenance (Fig. 3C, D). Consistent with RPGRIP1L knockdown, IFT88 depletion also resulted in a substantial reduction in lumen development compared to controls (Fig. 3F), reinforcing that functional primary cilia are required for lumenogenesis.

**Figure 3.**
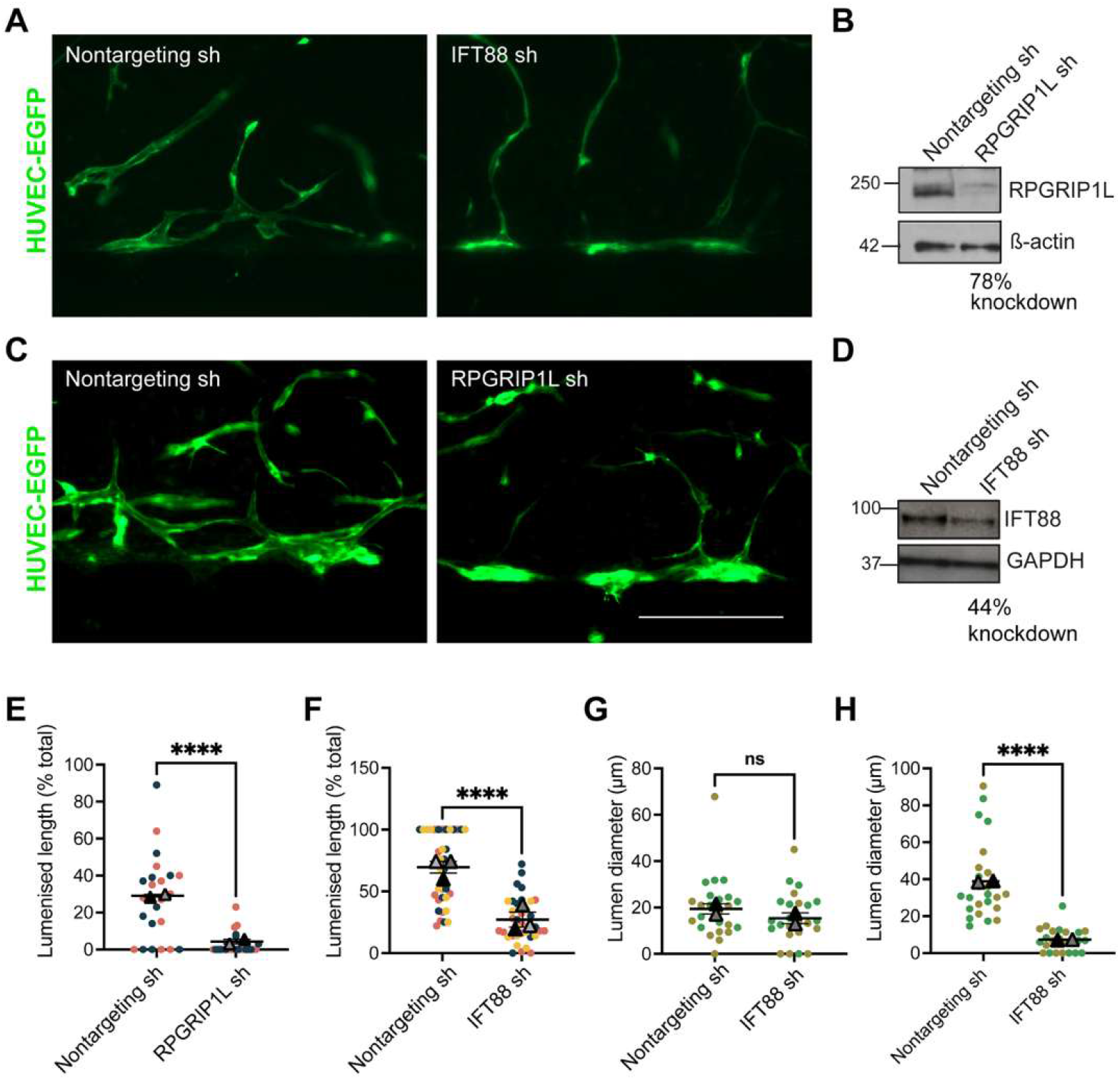
Ciliary proteins IFT88 or RPGRIP1L are necessary for lumen development under conditions of fluid flow. **(A)** Epifluorescence images of lumen formation on day 14 in HUVECEGFP cells expressing either non-targeting shRNA (Nontargeting sh) or RPGRIP1L shRNA (RPGRIP1L sh). shRNA expression was induced with doxycycline (2 µg/ml) between days 8–14. Scale bar, 400 µm. **(B)** Western blot confirming RPGRIP1L knockdown in a HUVEC monolayer after 72 h of doxycycline treatment. **(C)** Epifluorescence images of lumen formation on day 14 following induction of either non-targeting or IFT88 shRNA with doxycycline (2 µg/ml) between days 8–14 in HUVECEGFP cells. Scale bar, 400 µm. **(D)** Western blot confirming IFT88 knockdown in a HUVEC monolayer after 72 h of doxycycline treatment. **(E)** Quantification of lumenised length (% of total) for the experiment shown in (A). Data are presented as mean ± SEM from two independent experiments (n = 24 channel squares per condition). **(F)** Quantification of lumenised length (% of total) for the experiment shown in (C). Data are mean ± SEM from three independent experiments (n = 36 channel squares per condition). **(G)** Lumen diameter quantification before doxycycline induction (day 8) in control and IFT88 knockdown cells. **(H)** Lumen diameter quantification after doxycycline induction (day 14) in control and IFT88 knockdown cells. **(E, G)** Data are presented as mean ± SEM from two independent experiments (n = 24 channel squares). Statistical significance in all experiments was determined using a paired two-tailed Student’s t-test (ns, non-significant; ****, p < 0.0001).

The inducible nature of this system enabled temporal dissection of cilia function. Before doxycycline induction (day 8), control and IFT88-knockdown cells formed lumens with comparable diameters (Fig. 3G). However, following induction, lumen expansion was markedly impaired in the knockdown group by day 14 (Fig. 3H). These data demonstrate that primary cilia are essential not only for the initiation of lumen formation but also for its subsequent progression and maintenance.

### ROCK signalling negatively regulates ciliogenesis through LIMK-dependent actin stabilisation

We next investigated whether ROCK signalling regulates ciliogenesis. In hTERT-RPE1 epithelial cells, a well-established model for ciliogenesis, pharmacological ROCK inhibition induced dose-dependent cilia elongation, with an approximate 20% increase at 10 µM Y27632 and 50% at 50 µM (Fig. 4A, B). Given that ROCK modulates actin dynamics via LIM kinase (LIMK), we asked whether this pathway underlies its effect on cilia. Consistent with this model, inhibition of LIMK (using BMS-5) or actin depolymerisation (using latrunculin B) also significantly increased ciliary length in hTERT-RPE1 cells (Fig. 4C, D), suggesting that ROCK signalling restricts cilia elongation by stabilising the actin cytoskeleton through LIMK.

**Figure 4.**
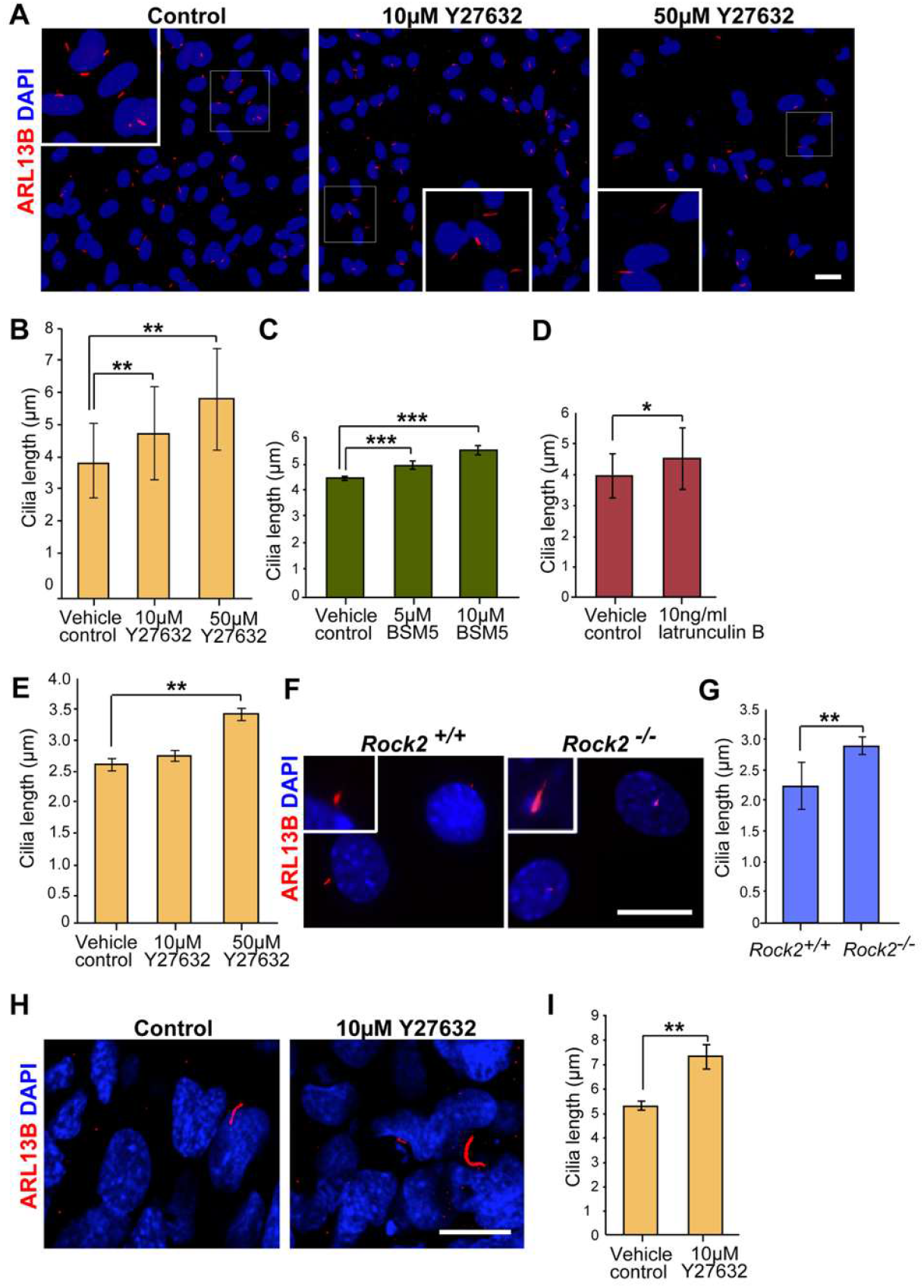
ROCK inhibition or *Rock2* genetic deletion deregulates cilia development. **(A)** Representative immunofluorescence images of ARL13Bstained cilia in serum starved hTERTRPEl cells treated with DMSO (control) or Y27632 (10 pM, 50 pM) for 48 h. Zoomed insets show individual cilia. Scale bar, 30 pm. **(B)** Quantification of cilia length in hTERTRPEl cells from (A). Data are mean ± SEM from four independent experiments (n > 100 cells per condition). **(C)** Cilia length quantification in hTERTRPEl cells treated with the LIMK inhibitor BMS5 (5 pM, 10 pM) for 48 h. Data are mean ± SEM from three independent experiments (n > 50 cells per condition). **(D)** Cilia length quantification in hTERTRPE1 cells treated with latrunculin B (0.01 pg/pl) for 48 h. Data are mean ± SEM from three independent experiments (n > 50 cells per condition). **(E)** Cilia length quantification in wildtype MEFs treated with Y27632 (10 pM, 50 pM) for 48 h. Data are mean ± SEM from three independent experiments (n > 50 cells per condition). **(F)** Representative immunofluorescence images of cilia in wildtype (WT) and Rock2/MEFs after 48 h serum starvation. **(G)** Quantification of average cilia length in MEFs shown in (F). Data are mean ± SEM (n > 50 cells per condition). **(H)** Representative immunofluorescence images of cilia in hCMEC treated with Y27632 (10 pM) for 48 h. **(I)** Quantification of average cilia length in ciliated hCMEC from (H). Data are mean ± SEM from three independent experiments (n > 30 cells per condition). Statistical significance was determined using a two-tailed Student’s *t* test (*, p < 0.05; **, p < 0.01; ***, p < 0.001)

To determine whether the elongation phenotype observed with Y27632 was specifically due to ROCK2 inhibition, we used wild-type and *Rock2*^-/-^ mouse embryonic fibroblasts (MEFs). Although MEF cilia were shorter (2–3 µm) than those in RPE1 cells (3–20 µm), wild-type MEFs exhibited a significant increase in ciliary length upon Y27632 treatment (Fig. 4E). Importantly, untreated *Rock2*^-/-^ MEFs displayed significantly longer cilia than wild-type controls (Fig. 4F, G), phenocopying the pharmacological inhibition.

Finally, we tested the effect of ROCK inhibition in human cerebral microvascular endothelial cells (hCMECs). Despite a low basal ciliation rate, Y27632 treatment significantly increased cilia length in these cells (Fig. 4H, I), confirming that ROCK negatively regulates ciliogenesis across multiple cell types, including endothelium.

### ROCK2 deficiency impairs blood vessel lumen formation and causes haemorrhages in mouse embryos

Previous studies demonstrated that global deletion of mouse *Rock2* results in embryonic lethality, with fewer than 10% of *Rock2* knockout (KO) embryos surviving beyond embryonic day (E) 13.5.^24^ Surviving mice are growth-retarded but develop normally and remain fertile. Embryonic lethality was attributed to placental dysfunction and intrauterine growth restriction prior to foetal death.

Consistent with published observations^24^, in our analysis *Rock2*^-/-^embryos harvested at E13.5 appeared smaller than their wild-type littermates and exhibited macroscopic haemorrhages in the head, neck, and trunk regions, with occasional pooled blood in the paws and feet (Fig. 5A).

Some *Rock2*^-/-^ embryos also displayed omphalocele (Fig. 5A). These phenotypic features prompted further investigation into the role of ROCK2 in vascular lumen formation and ciliary regulation during embryonic development.

Immunohistochemistry confirmed the absence of ROCK2 protein and revealed severe pericardial haemorrhage associated with a fractured and discontinuous epicardium and an enlarged aorta (Fig. 5B,C). Defects in vascular integrity were also evident in superficial brain blood vessels, where blood cells were found outside vessels exclusively in *Rock2* KO embryos, indicating systemic loss of vascular integrity (Fig. 5D).

Given the haemorrhagic phenotypes, we assessed lumen formation in the embryonic brain. Transverse sections at the level of the interventricular foramen of Monro were stained for CD31, and parenchymal vessels within the cerebral hemispheres were quantified (Fig. 5E). While the total number of CD31-positive vessels was unchanged in *Rock2*^-/-^ embryos (Fig. 5F), the number of vessels with open, perfused lumens was significantly reduced (Fig. 5G). Furthermore, the lumens that did form were significantly larger in diameter than those in wild-type littermates (Fig. 5H).

Together, these data demonstrate that ROCK2 is essential for proper lumen formation and the maintenance of vascular integrity during embryonic development.

### ROCK2 regulates endothelial ciliogenesis and cilia orientation in the developing brain

Given our *in vitro* finding that ROCK2 regulates cilia length, we next asked whether it controls endothelial ciliogenesis *in vivo*, particularly in the embryonic brain where cilia are known to play critical roles during vascular development^25,26^ (Fig. 6A). We quantified endothelial cilia in the brain parenchyma of E13.5 wild-type (WT) and *Rock2* KO mouse embryos (Fig. 6A, B). Tissue sections were immunostained for ARL13B, and cilia associated with blood vessels were counted to assess changes in both abundance and spatial orientation (Fig. 6B). To characterise how cilia are spatially organised within the vessel wall, we categorised their orientation relative to the lumen: Luminal (projecting into the lumen), Internal (within the endothelial layer), or External (projecting into the parenchyma) (Fig. 6C).

**Figure 6.**
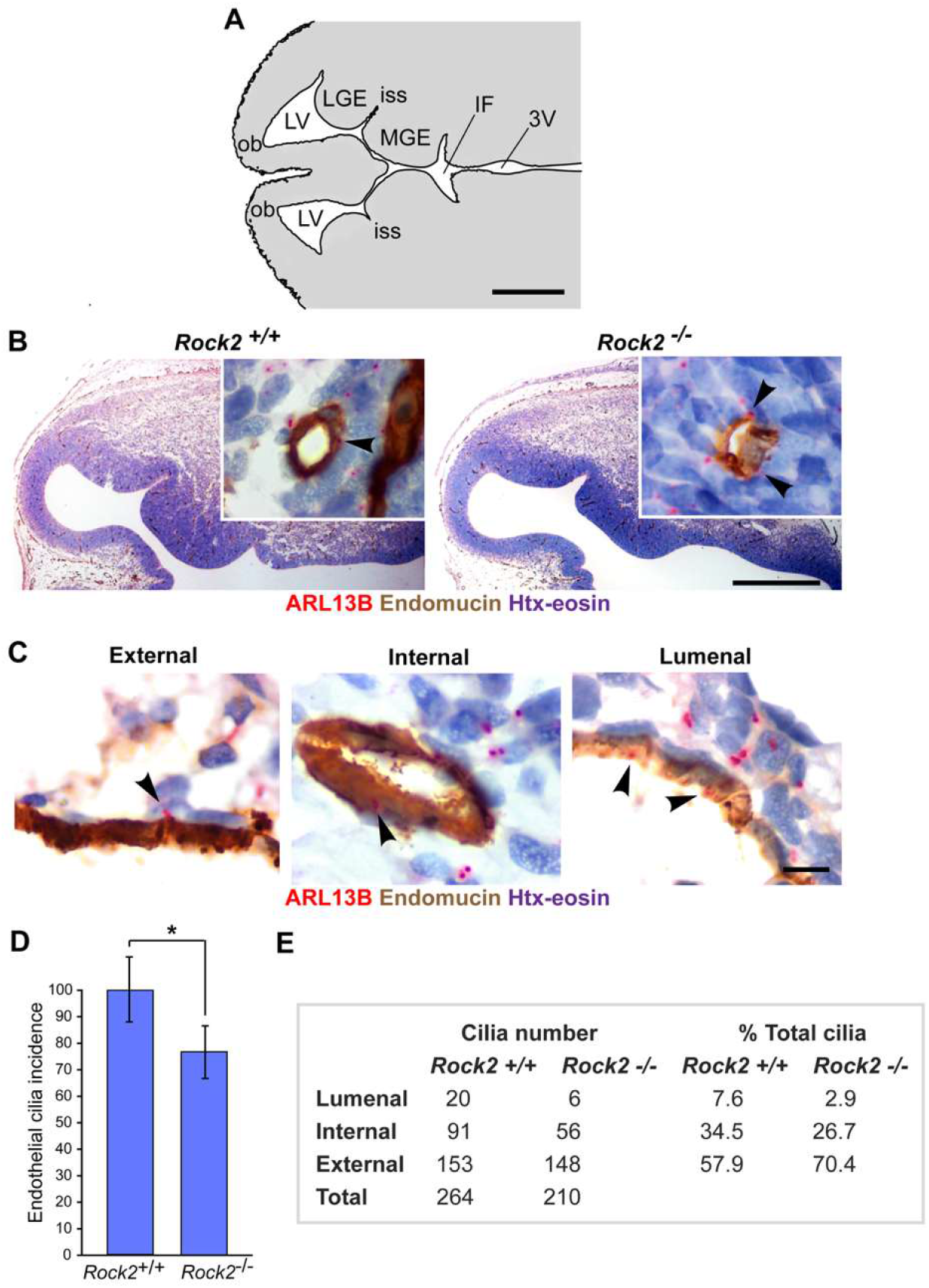
*Rock2* genetic deletion disrupts vascular cilia development and positioning *in vivo*. **(A)** Schematic of a transverse mouse brain section at the level of the interventricular foramen of Monro (the channel connecting the lateral ventricles to the third ventricle). 3V, third ventricle; IF, interventricular foramen; iss, intermediate subpallial sulcus; LGE, lateral ganglionic eminence; LV, lateral ventricle; MGE, medial ganglionic eminence; ob, olfactory bulb. Scale bar, 1 mm. **(B)** Representative images of H&E sections immunolabeled for endomucin (brown, vasculature) and ARL13B (red, cilia) in Rock2^+/+^ and Rock2^−/−^ E13.5 embryos. Scale bar, 200 µm. Zoomed insets show individual vessels. Scale bar,100 µm. **(C)** High magnification representative images illustrating ciliary positioning categories: Luminal (projecting into the lumen), Internal (within the endothelial layer), and External (projecting into the parenchyma). Arrowheads indicate primary cilia. Scale bar, 10 µm. **(D)** Quantification of total endothelial cilia counts in the brain parenchyma of Rock2^+/+^ and Rock2^−/−^ embryos. Data are mean ± SEM; n ≥ 50 vessels per embryo (Rock2^+/+^, 3; Rock2^−/−^, 3). **(E)** Quantification of the percentage of cilia in each positional category (Luminal, Internal, External) in Rock2^+/+^ and Rock2^−/−^ embryos. Table summarising Endothelial cilia counts in parenchymal blood vessels of Rock2^+/+^ and Rock2^−/−^ embryos. n = number of embryos (Rock2^+/+^, 3; Rock2^−/−^, 3). Statistical significance was determined using a two-tailed Student’s *t* test (*, p < 0.05; ***, p < 0.001).

The analysis revealed a ∼20% reduction in total endothelial cilia number in the blood vessels of Rock2 KO embryos compared to WT controls (Fig. 6D). Strikingly, we observed a profound positional defect: in *Rock2* KO embryos, the proportion of internal/luminal cilia dropped from 42.1% to 29.6%, while externally oriented cilia increased from 57.9% to 70.4% (Fig. 6E).

Thus, ROCK2 is essential not only for establishing normal cilia numbers but also for their precise orientation within the vessel wall, implicating it as a key regulator of endothelial cilia architecture during brain vascular development.

## Discussion

The primary cilium is a mechanosensory organelle that responds to extracellular inputs, playing diverse roles in signal transduction and developmental signalling.^27^ In vascular endothelial cells, primary cilia are thought to sense and transduce fluid shear into biochemical signals, supported by the observation that each endothelial cell possesses a single apical cilium containing polycystin-1 and IFT88.^12^ In this study, we show that cilia ablation through knockdown of IFT88 or RPGRIP1L, or cilia deregulation through ROCK inhibition, impairs lumen formation and expansion in a perfused microfluidic model. Using a *Rock2* mouse knockout model, we further demonstrate that loss of ROCK2 reduces endothelial cilia incidence and alters their positioning relative to the vessel lumen, phenotypes associated with disrupted lumen formation and haemorrhage in the embryonic brain. These findings uncover a novel co-regulatory axis between ROCK signalling and primary cilia essential for vascular lumen development.

Haemodynamic forces—shear stress and pressure-derived stretch—are key drivers of vascular morphogenesis.^11^ During lumen expansion, blood pressure drives inverse blebbing,^4^ while shear stress induces EC polarisation, junctional remodelling, and transcriptional responses.^5,28^ The primary cilium is a critical mechanosensor for low flow, and its dysfunction phenocopies flow-loss defects.^14^ Our work extends these findings by positioning ROCK2 upstream of ciliary mechanosensation, regulating cilia abundance and orientation. Flow-dependent vessel calibre regulation involves multiple mechanisms, including EC shape changes mediated by endoglin^29^ and YAP nuclear translocation under high shear.^30^ Our observation that ROCK2 loss alters lumen diameter (Fig. 5H) suggests that ROCK2 may integrate with these pathways. Mechanistically, we demonstrate that ROCK disruption leads to cilia deregulation via LIM-kinase inhibition and dysregulated actin polymerisation, positioning ROCK2 as a key upstream regulator of endothelial cilia.

An emerging link between ciliary function and the cytoskeleton is supported by findings that ROCK2 influences cilia formation through actomyosin contractility.^31^ Several zebrafish and mouse ciliopathy models display vascular anomalies—haemorrhage, vessel dilatation, and defective branching—with mutations in ciliary genes such as IFT88, KIF3A, and TTC21B impairing endothelial cilia and disrupting developmental angiogenesis.^15,32,33^ Importantly, Kallakuri et al.^15^ demonstrated that endothelial-specific re-expression of IFT genes rescues intracranial haemorrhage in zebrafish ciliary mutants, establishing an endothelial-autonomous role for cilia in vascular integrity. In humans, ciliary dysfunction underlies autosomal dominant polycystic kidney disease (ADPKD), associated with increased cardiovascular complications including hypertension, intracranial aneurysms, and myocardial infarction, underscoring the clinical importance of endothelial cilia in vascular health.^34,35,36^ Consistent with recent findings from endothelial-specific Rock1/2 deletion,^37^ our *Rock2* global knockout embryos exhibited haemorrhages and defective lumen formation, supporting a cell-autonomous role for endothelial ROCK2 in vascular integrity.

Microfluidic devices enable exposure of cells to controlled flow and shear stress, recapitulating key aspects of in vivo microenvironments.^19^ The 5-pillar device used in this study supports the development of hierarchical lumens and capillary-like tubes (10–15 µm) that approach the dimensions of cerebral capillaries. While a 3-pillar version has been described^20^, that design produces different flow profiles and larger vessels (>50 µm) representative of precapillary arterioles. We employed this platform to monitor lumen development and cilia disruption under physiologically relevant flow conditions, allowing real-time observation, pharmacological perturbation, and genetic manipulation of developing lumens.

Collectively, our findings support a model in which ECs integrate multiple mechanical inputs—shear stress sensed by cilia and pressure-derived stretch—through ROCK2-dependent cytoskeletal regulation to coordinate lumen formation, expansion, and stabilisation. This framework aligns with emerging views that EC mechanosensation is a network of overlapping sensors and effectors,^11^ unveiling a previously unrecognised co-regulatory axis essential for vascular lumen formation.

## Acknowledgements

This work was supported by the British Heart Foundation (FS/18/32/33557 to G.M. and C.A.J.) and the Medical Research Council (MR/M000532/1 to C.A.J.). S.Z.M. was funded by the BHF grant. G.G. was supported by a University of Leeds Research Scholarship. The authors thank M. Olson (University of Toronto) and G. Whyte (Heriot-Watt University) for discussions.

## Author contributions

**Formatted:** Font: Italic

**Formatted:** Font: Italic

**Formatted:** Font: Italic

G.M. and C.A.J. conceived and supervised the study and secured the funding. S.Z.M. performed and analysed the microfluidic experiments under flow. A.M. generated the *Rock2*^-/-^ mice, set up timed matings, and generated *Rock2*^-/-^ MEFs. G.G. performed cilia visualisation and analyses and characterised the *Rock2*^-/-^ mice. M.D.B. and S.A.P. designed the microfluidic devices. S.Z.M. fabricated the microfluidic devices and performed the microfluidic experiments. All authors contributed to the writing and revision of the manuscript and approved the final version.

## Data availability

Data for this article are available at the White Rose eTheses Online repository oai:etheses.whiterose.ac.uk:32116 and oai:etheses.whiterose.ac.uk:17044

## References

1. Lammert E, Cleaver O, Melton D. Induction of pancreatic differentiation by signals from blood vessels. Science. 2012;338(6106):458–462.

2. Sigurbjornsdottir S, Mathew R, Leptin M. Molecular mechanisms of de novo lumen formation. Nat Rev Mol Cell Biol. 2014;15(10):665–676.

3. Strilić B, Kucera T, Eglinger J, et al. The molecular basis of vascular lumen formation in the developing mouse aorta. Dev Cell. 2009;17(4):505–515.

4. Gebala V, Collins R, Geudens I, Phng LK, Gerhardt H. Blood flow drives lumen formation by inverse membrane blebbing during angiogenesis. Dev Cell. 2016;37(3):201–213.

5. Franco CA, Jones ML, Bernabeu MO, et al. Dynamic endothelial cell rearrangements drive developmental vessel regression. PLoS Biol. 2016;14(6):e1002475.

6. Kwon HB, Wang S, Helker CS, et al. In vivo modulation of endothelial polarization by apelin signalling. Nat Commun. 2016;7:11847.

7. Bryan BA, D’Amore PA. What tangled webs they weave: Rho-GTPase control of angiogenesis. Cell Mol Life Sci. 2007;64(16):2053–2065.

8. Barry AK, Wang N, Leckband DE. Local RhoA activation induces formin-dependent apical domains and lumenogenesis. Mol Biol Cell. 2016;27(18):2788–2799.

9. Hartmann S, Ridley AJ, Lutz S. The function of Rho-associated kinases ROCK1 and ROCK2 in the pathogenesis of cardiovascular disease. Front Pharmacol. 2015;6:276.

10. Lindsey SE, Butcher JT, Yalcin HC. Mechanical regulation of cardiac development. Front Physiol. 2018;9:1560.

11. Campinho P, Vilfan A, Vermot J. Blood flow forces in shaping the vascular system: a focus on endothelial cell behavior. Front Physiol. 2020;11:552.

12. Nauli SM, Kawanabe Y, Kaminski JJ, et al. Endothelial cilia are fluid shear sensors that regulate blood pressure. J Clin Invest. 2008;118(3):949–956.

13. Egorova AD, Khedoe PP, Goumans MJ, et al. Lack of primary cilia primes shear-induced endothelial-to-mesenchymal transition. Circ Res. 2012;111(10):1306–1315.

14. Goetz JG, Steed E, Ferreira RR, et al. Endothelial cilia mediate low flow sensing during zebrafish vascular development. Cell Rep. 2014;6(5):799–808.

15. Kallakuri S, Yu JA, Li R, et al. Endothelial cilia are essential for vascular integrity in zebrafish. Development. 2015;142(6):1035–1045.

16. Kim J, Lee JE, Heynen-Genel S, et al. Functional genomic screen for modulators of ciliogenesis and cilium length. Nature. 2010;464(7291):1048–1051.

17. Kim S, Lee EJ, Kim J. Cell-cycle-dependent oscillation of the ciliary length. Cilia. 2015;4(Suppl 1):P72.

18. Hetheridge C, Mavria G, Mellor H. Uses of the in vitro endothelial-fibroblast organotypic co-culture assay in angiogenesis research. Biochem Soc Trans. 2011;39:1597–1600.

19. Moya ML, Hsu YH, Lee AP, Hughes CC, George SC. In vitro perfused human capillary networks. Tissue Eng Part C Methods. 2013;19:730–737.

20. Bourn MD, Zahed Mohajerani S, Mavria G, Ingram N, Coletta PL, Evans SD, et al. Tumour associated vasculature-on-a-chip for the evaluation of microbubble-mediated delivery of targeted liposomes. Lab Chip. 2023;23:1674–1693.

21. Kümper S, Mardakheh FK, McCarthy A, Yeo M, Stamp GW, Paul A, et al. Rho-associated kinase (ROCK) function is essential for cell cycle progression, senescence

22. Vion AC, Alt S, Klaus-Bergmann A, Szymborska A, Zheng T, Perovic T, et al. Primary cilia sensitize endothelial cells to BMP and prevent excessive vascular regression. J Cell Biol. 2018;217:1651–1665.

23. Ishikawa H, Marshall WF. Ciliogenesis: building the cell’s antenna. Nat Rev Mol Cell Biol. 2011;12:222–234.

24. Thumkeo D, Keel J, Ishizaki T, Hirose M, Nonomura K, Oshima H, et al. Targeted disruption of the mouse rho-associated kinase 2 gene results in intrauterine growth retardation and fetal death. Mol Cell Biol. 2003;23(14):5043–5055.

25. Eisa-Beygi S, Benslimane FM, El-Rass S, Prabhudesai S, Abdelrasoul MKA, Simpson PM, et al. Characterization of endothelial cilia distribution during cerebral-vascular development in zebrafish (Danio rerio). Arterioscler Thromb Vasc Biol. 2018;38:2806–2818.

26. Prabhudesai S, Thirugnanam K, Song X, Yang H, Errede M, Girolamo F, et al. Brain vascular stability relies on PAK2-cilia-PDGF-BB-HSPGs on basolateral side of endothelium. Life Sci Alliance. 2026;9:e202503021.

27. Smith CEL, Hayward V, Wilson R. Primary cilia in development and disease. Development. 2020;147(15):dev188367.

28. Baeyens N, Mulligan-Kehoe MJ, Corti F, et al. Syndecan 4 is required for endothelial cell alignment in flow. J Cell Biol. 2015;210(6):937–951.

29. Sugden WW, Meissner R, Aegerter-Wilmsen T, et al. Endoglin controls blood vessel diameter through endothelial cell shape changes in response to haemodynamic cues. Nat Cell Biol. 2017;19(6):653–665.

30. Nakajima H, Yamamoto K, Agarwala S, Terai K, Fukui H, Matsuda M. Flow-dependent endothelial YAP translocation contributes to vascular maintenance. Dev Cell. 2017;40(1):6–11.

31. Smith CS, Kim CY, Li R, et al. ROCK2 regulates ciliogenesis through actomyosin contractility. J Cell Sci. 2025;138(4):jcs261234.

32. Elworthy S, Hargrave M, Knight R, Muiño-Blanco T, Kelsh RN. Ciliary genes are required for vascular development in zebrafish. Dis Model Mech. 2019;12(5):dmm038976.

33. Ma M, Gallagher AR, Somlo S. Ciliary mechanisms in cystic kidney disease. J Clin Invest. 2020;130(7):3370–3381.

34. Lin SL, Chiang HY, Chen YM, et al. Cardiovascular complications in autosomal dominant polycystic kidney disease. J Am Soc Nephrol. 2003;14(9):2293–2299.

35. Pietrzak-Nowacka M, Nowak M, Krzanowski M. Cardiovascular manifestations in patients with autosomal dominant polycystic kidney disease. Medicina. 2024;60(3):412.

36. Ahmed S, Ali SM, Farooq S. Hypertension and cardiovascular risk in ADPKD: a systematic review. J Clin Med. 2025;14(2):456.

37. Lange M, Sinha A, Wanner P, et al. Endothelial ROCK1 and ROCK2 regulate vascular lumen formation and integrity. Circ Res. 2026;138(5):e45–e60.

